# Taxonomic assessment of two wild house mouse subspecies using whole-genome sequencing

**DOI:** 10.1101/2022.06.28.497812

**Authors:** Raman Akinyanju Lawal, Verity L. Mathis, Mary E. Barter, Jeremy R. Charette, Alexis Garretson, Beth L. Dumont

## Abstract

House mice (*Mus musculus*) are comprised of three primary subspecies. A large number of secondary subspecies have also been suggested on the basis of divergent morphology and molecular variation at limited numbers of markers. While the phylogenetic relationships among the primary *M. musculus* subspecies are well-defined, the relationships among the secondary subspecies, and their relationships to primary subspecies, remain poorly understood. Here, we integrate *de novo* genome sequencing of museum-stored specimens of house mice from one secondary subspecies (*M. m. bactrianus*) and publicly available genome sequences of house mice previously characterized as *M. m. helgolandicus*, with whole-genome sequences from diverse representatives of the three primary house mouse subspecies. We show that mice assigned to the secondary *M. m. bactrianus* and *M. m. helgolandicus* subspecies are not genetically differentiated from the contemporary *M. m. castaneus* and *M. m. domesticus*, respectively. Overall, our work suggests that the *M. m. bactrianus* and *M. m. helgolandicus* subspecies are not well-justified taxonomic entities, emphasizing the importance of leveraging whole-genome sequence data to inform subspecies designations. Additionally, our investigation has established experimental procedures for generating whole-genome sequences from air-dried museum specimens and provides key genomic resources to inform future genomic investigations of wild mouse diversity.

## Introduction

House mice (*M. musculus*) are the premiere mammalian model system for biomedical research and an important natural model system for ecological and evolutionary studies ^1^. House mice emerged from an ancestral population in the Indian subcontinent less than 3 million years ago ^2,3^ and subsequently expanded out of this ancestral region, giving rise to three primary subspecies ^4–7^. *M. m. domesticus* is native to Western Europe, *M. m. musculus* is present across Eastern Europe and Siberia, and *M. m. castaneus* expands across South and Southeast Asia. Aided by human dispersal in recent history, house mice have subsequently expanded their footprint outside of these native ranges, colonizing all major continents except Antarctica and many remote oceanic islands.

Beyond these three primary subspecies, a number of secondary house mouse subspecies have also been suggested on the basis of distinct morphology ^8–10^ and surveys of limited numbers of molecular markers ^5,10–15^. For example, mice from Yemen and Madagascar have been assigned to *M. m. gentilulus* due to their small size and distinct mitochondrial lineage ^11,13^. Mice from Heligoland, a small German archipelago island in the North Sea, have been characterized as *M. m. helgolandicus* due to their unique skull morphology, distinct mitochondrial D-loop haplotype, and allelic variation at four nuclear loci ^10,16^. Similarly, mitochondrial sequence analysis has been used to assign house mice from Pakistan to the subspecies *M. m. bactrianus* ^15,17^. *M. m. musculus* and *M. m. castaneus* naturally hybridize where their ranges overlap in Japan, and these hybrids have been designated as a distinct subspecies, *M. m. molossinus* ^18^. At least seven other secondary subspecies of *M. musculus* have been named, including: *M. m. albula, M. m. brevirostris, M. m. gansuensis, M. m. homourus, M. m. isatissus, M. m. wagneri*, and *M. m. gansuensis* ^19^.

Prior studies have leveraged powerful genomic approaches to investigate the evolutionary relationships between the three primary house mouse subspecies, establishing a sister relationship between *M. m. castaneus* and *M. m. musculus* ^20–23^. In contrast, the phylogenetic relationships among secondary house mouse subspecies, including their relationships to the primary house mouse subspecies, remain poorly understood. Virtually all secondary subspecies assignments have been informed by sparse molecular data, begging the question of whether distinct subspecies labels are truly warranted. In particular, the absence of whole-genome sequence data has made it impossible to determine whether specific secondary subspecies designations represent premature conclusions based on chance sampling of population-private alleles or are justified based on genome-wide patterns of divergence.

Here, we combine published whole genome sequencing data ^24–26^ with *de novo* genome sequencing of a strategic set of museum-preserved specimens to address the validity of two secondary house mouse subspecies designations - *M. m. bactrianus* and *M. m. helgolandicus*. Our investigations offer a conclusive resolution to the taxonomic status of mice from these subspecies, arguing that these taxonomic assignments are not well justified based on genomic data.

## Results and Discussion

### Whole-genome sequencing of museum samples of wild-caught mice from Pakistan

We whole-genome sequenced 14 ~46-year-old museum-preserved specimens of *M. m. bactrianus* from Pakistan (PAK) to 17x – 45x coverage (Supplementary Fig. S1a). These museum samples were collected across several counties in Pakistan and maintained by the Florida Museum of Natural History (Fig. 1). This geographic sample region overlaps with the presumed ancestral home of house mice ^5^. On average, ~99% of sequenced reads from each of these mouse genomes were mapped to the mm10 reference (Supplementary Fig. S1b), suggesting little exogenous contamination of our samples. The proportion of bases supported by at least 5 reads ranges from ~66% to 96% across samples (Supplementary Fig. S1c).

**Figure 1.**
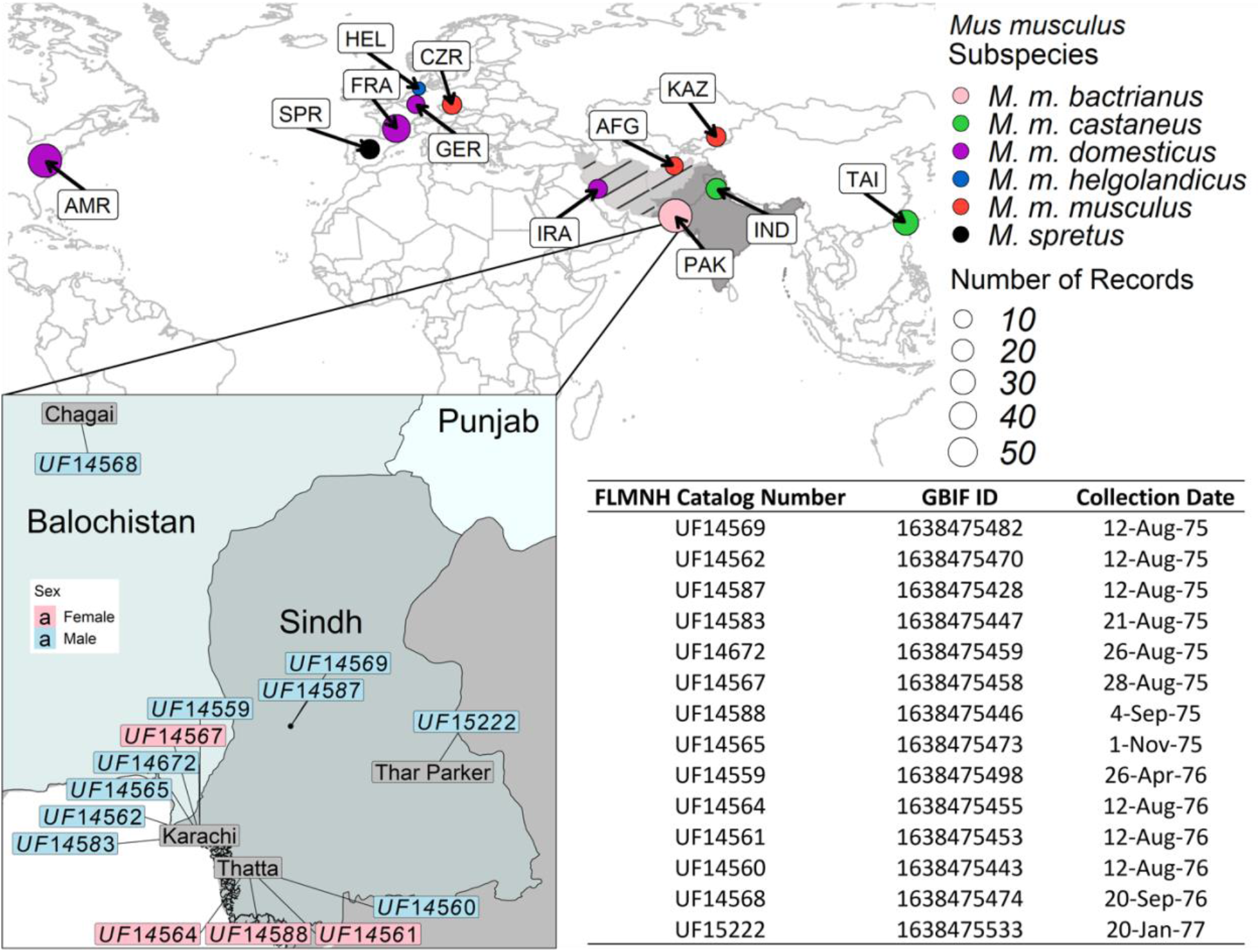
Map showing the approximate geographic sampling locations of house mouse populations profiled by whole-genome sequencing. The house mouse ancestral region extends from India “IND” and Pakistan “PAK” (dark grey area) and may include the broader region from Iran “IRA” and Afghanistan “AFG” (light grey area with black markings). All genome sequences except for PAK were retrieved from public databases ^24–26^. The genome sequences also include populations from America (AMR), France (FRA), Germany (GER), Heligoland (HEL), Kazakhstan (KAZ), and Czech Republic (CZR), Taiwan (TAI), and *M. spretus* from Spain (SPR). The enlarged plot inset shows the locations of PAK samples collected across four counties in the Sindh region of Pakistan. Florida Museum of Natural History (FMNH) Catalog numbers, Global Biodiversity Information Facility (GBIF) identifiers ^30^, and collection dates for each PAK sample are presented in the table.

Ancient and museum-stored specimens are known to accumulate C → T and A → G mutations due to post-mortem DNA deamination ^27^. This spontaneous damage can lead to sequence biases if not properly accounted for. We determined that, across these PAK genomes, the deamination rate for nucleotide misincorporation is ~0.1%, with nearly all these damages confined to the first and last 5 bp of a given read (Supplementary Fig. S2). By comparison, the frequency of post-mortem DNA damage in a sample of ancient (~13k years old) stickleback fish was 30% ^28^ and is typically around 50% for Neanderthal DNA ^29^. Although there is minimal post-mortem DNA in our sequenced PAK genomes, we took the conservative approach of hard clipping the terminal 5 bp on both the 5’ and 3’ ends of every read to eliminate potential artifacts (Supplementary Fig. S1d).

After these quality control steps and applying basic hard filters to eliminate low-quality calls (see Methods), we identified a total of 81,338,251 autosomal SNPs and 8,802,753 short indels across the 14 PAK genomes. This corresponds to ~1 variant every 30 bases relative to the mm10 reference genome.

### Re-evaluating phylogenetic relationships among *M. musculus* subspecies

We combined the 14 PAK genome sequences with 3 previously published genome sequences of *M*. *m. helgolandicus* from Heligoland (HEL) and 152 publicly available whole-genome sequences from wild-caught mice from multiple populations for each *M. m. castaneus* (CAS), *M. m. musculus* (MUS), and *M. m. domesticus* (DOM) ^24–26^. Using this comprehensive set of 169 whole genome sequences, we evaluated the genetic relationships among five putative subspecies using a multi-pronged approach.

First, we conducted a principal component analysis (Fig. 2a). The first two principal components (26.14% variance along PC1 and 18.21% of PC2) reveal just three discrete clusters (G1-G3) corresponding to the primary MUS, DOM, and CAS subspecies. Mice from the HEL population group within DOM, while PAK is nearly indistinguishable from CAS populations (Fig. 2a). Additional approaches based on average pairwise distance among individuals (Fig. 2b), allele sharing distance (Fig. 2c), and co-ancestry (Fig. 2d) also support three discrete phylogenetic units aligning to the three primary *M. musculus* subspecies.

**Figure 2:**
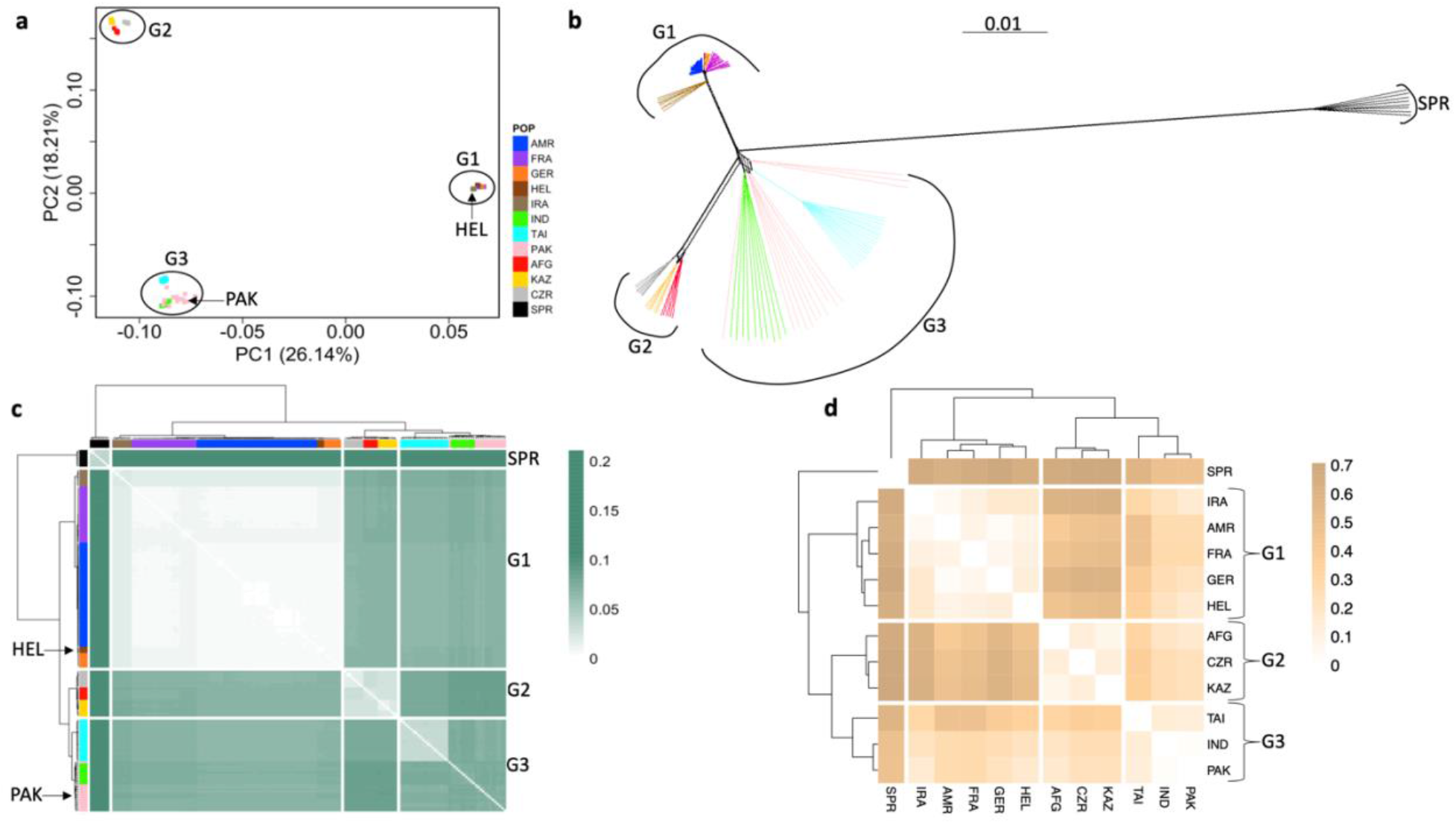
Genetic relationships among five putative *M. musculus* subspecies reveal three taxonomic groups (G1 – G3). Genetic relationships were assessed via (**a**) principal component analysis, (**b**) a phylogenetic tree constructed from a pairwise distance matrix, (**c**) allele sharing distance, and (**d**) co-ancestry based on FST. The color legend on the right side of panel (**a**) is also applicable to panels (**b**) and (**c**). G1 groups populations belonging to DOM subspecies: America (AMR), France (FRA), Germany (GER), Heligoland (HEL), and Iran (IRA). G2 groups populations of MUS: Afghanistan (AFG), Kazakhstan (KAZ), and Czech Republic (CZR). G3 groups CAS populations: India (IND), Taiwan (TAI), and Pakistan (PAK). *M. spretus* (SPR) is used as an outgroup.

Wild mice may experience substantial gene flow between populations and subspecies ^31^, which could obscure subspecies relationships. To evaluate the possible influence of gene flow on our inferred subspecies groupings, we used TreeMix to assess the robustness of our findings under multiple distinct migration models ^32^. We recovered identical genealogical relationships both in the absence of gene flow and under different migration scenarios (*p* < 1×10^-300^; Fig. 3). Importantly, no tested migration model altered the composition of the three core subspecies groups or offered support for the *M. m. helgolandicus* or *M. m. bactrianus* subspecies designations.

**Figure 3:**
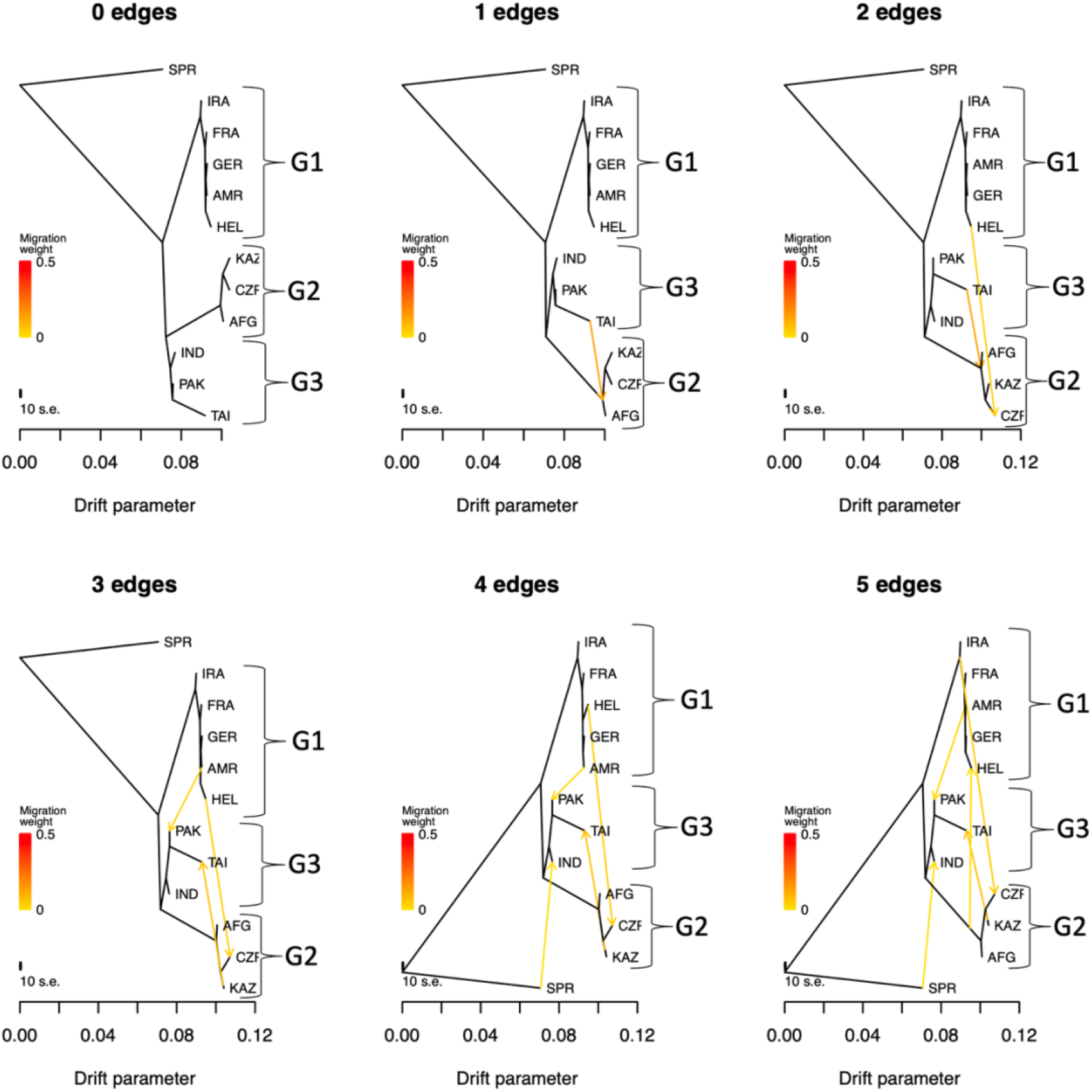
Phylogenetic relationships from TreeMix between house mouse populations. Treemix modeling the effect of different numbers of populations experiencing gene flow and different pairwise migration rates between populations. Zero edges correspond to the absence of gene flow. Under all surveyed scenarios, HEL is embedded within the taxonomic group G1 (*M. m. domesticus*) and PAK in G3 (corresponding to *M. m. castaneus* populations). *M. spretus* (SPR) was used as an outgroup.

We find no compelling evidence to justify a unique subspecies designation for mice from HEL or PAK. These conclusions contradict earlier findings based on limited numbers of genomic markers ^10^ or single loci ^15^, underscoring the power and importance of leveraging whole-genome data to inform subspecies designations.

Our investigations suggest that PAK and HEL are populations of *M. m. castaneus* and *M. m. domesticus*, respectively. Combining these populations with their modified subspecies groupings, we next partitioned the genome into 13,122 blocks, constructed phylogenetic trees from each partition, and estimated the percentage of the genome supporting each of the three possible topologies relating the three primary *M. musculus* subspecies. Overall, the topology placing CAS and MUS as sister is the most abundant in the genome, capturing a total weight of 34.20% (autosomes), 36.70% (X chromosome), and 44.20% (MT) (Fig. 4). This finding validates prior conclusions about house mouse subspecies phylogenetic relationships based on representative inbred strain genomes ^20,21^ and smaller samples of wild mice ^22^. Notably, our use of a broader set of wild house mouse genomes may provide more accurate estimates of the proportional representation of each topology and the extent of incomplete lineage sorting across the *M. musculus* genome.

**Figure 4:**
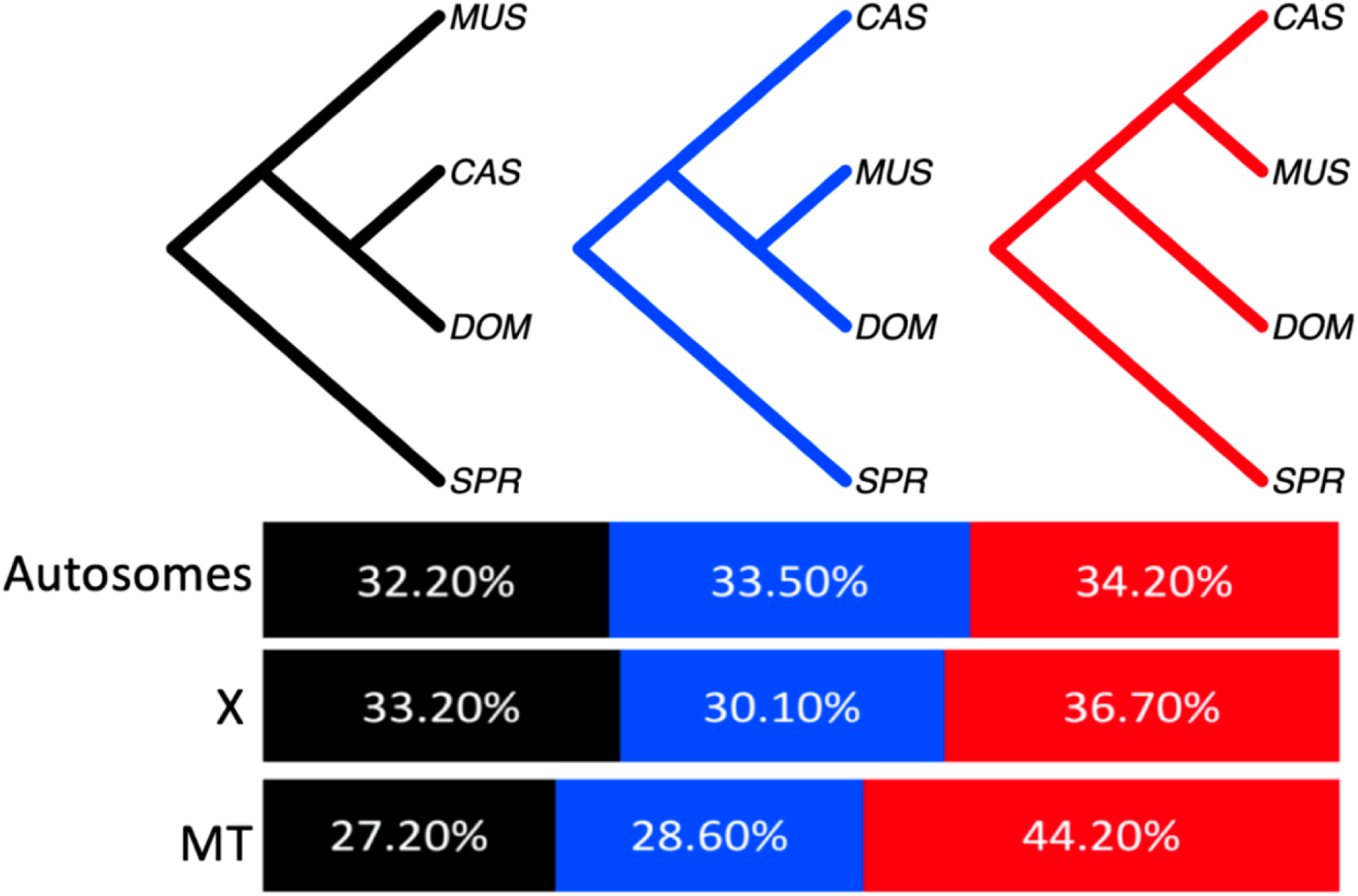
The percentage of the autosomal, X, and mitochondrial (MT) genome supporting each of the three possible topological relationships among the three primary house mouse subspecies. The topologies were derived from the construction of total of 13,122 phylogenetic trees across the genome. *M. m. castaneus* (CAS), *M. m. musculus* (MUS), *M. m. domesticus* (DOM), and outgroup *M. spretus* (SPR).

## Conclusions

Using both new and published whole-genome sequences from diverse wild house mice, we addressed support for two secondary *M. musculus* subspecies assignments: *M. m. bactrianus* and *M. m. helgolandicus*. We showed that mice from Pakistan previously assigned to *M. m. bactrianus* are genetically indistinguishable from *M. m. castaneus* mice. Similarly, mice assigned to the *M. m. helgolandicus* subspecies exhibit no meaningful genome-wide pattern of divergence from *M. m. domesticus*. While these subspecies may harbor distinct morphological adaptations ^10^, the adoption of a strict genetic species concept argues that *M. m. bactrianus* and *M. m. helgolandicus* are not well-justified taxonomic groups. Instead, mice previously assigned to these subspecies appear to capture population-level genetic and phenotypic diversity within the primary house mouse subspecies. Our work motivates additional genomic investigations into whether other secondary house mouse subspecies designations are warranted.

In addition to providing novel insights into house mouse taxonomy, our work yields newly sequenced house mouse genomes that will serve as key genetic resources for future investigations into wild mouse demographic history and diversity. In particular, genome sequencing of wild mice sampled from the cradle of *M. musculus* evolution will enable investigations into the impact of ancestral variation on contemporary patterns of global wild mouse diversity. Further, our work has established a robust experimental protocol and analytical pipeline for whole genome sequencing air-dried museum specimens. Broader application of this methodology to additional museum samples will offer a facile approach for strategically expanding genomic catalogs of wild mouse diversity.

## Materials and Methods

### Museum sample collection

We destructively sampled the skins of 14 *M. m. bactrianus* mice housed in the Florida Museum of Natural History Mammalogy Collection (http://specifyportal.flmnh.ufl.edu/mammals/). These specimens were collected and preserved between 1975 and 1977 across four counties in Pakistan ‘PAK’ (see Fig. 1 inset). Further details can be found in the GBIF.org (25 April 2022) GBIF occurrence download (https://doi.org/10.15468/dl.xuksm3) ^30^. Skin snips were obtained by removing a small (approx. 5 x 5 mm) section of skin from the ventral side of the study skin, sterilizing instruments in between samples.

### DNA extraction protocol for museum-stored samples

DNA was isolated from desiccated skin samples following a previously published protocol ^33^ with minor modifications. Briefly, the skin samples were scraped with a sterilized scalpel to remove possible contaminants. Samples were transferred to a 2 ml tube, washed three times with sterile water, three times with 70% ethanol, three times with sterile water, and then cut into small pieces. Samples were hydrated before digestion by incubating for 24 hours in 1 mL of TE (10mM Tris; 1mM EDTA, pH 7.6), washing with 70% ethanol and sterile water, and hydrating again in TE solution for a further 24 hours. Samples were digested in TNE solution (10 mM Tris HCl, pH 8; 400 mM NaCl; 2mM EDTA, pH 8.0) plus SDS 1% and Proteinase K (0.58 mg/ml final conc.) at 55 °C for 24-36 hours until the tissue was completely digested. The DNA was extracted with one volume of phenol:chloroform:isoamilic alcohol (25:24:1), rotated at 20 rpm for 10 minutes, and centrifuged for 10 minutes at 4000 rcf, after which the supernatant solution was transferred to another tube. The DNA was precipitated by adding two volumes of 100% ethanol, gently inverting the tube, and maintaining the solution at −20 °C for 16 h. The samples were centrifuged for 2 minutes at 3000 rcf before discarding the ethanol and resuspending the pellet in 50 μL of TE.

DNA sizing and concentration were assessed by TapeStation. DNA fragment sizes ranged from 76 bp to 431 bp. Only samples with DNA concentration > 63 ng/μl were used for genome sequencing.

### Genomic DNA library preparation

Whole-genome libraries were constructed using the KAPA HyperPrep Kit (Roche Sequencing and Life Science) according to the manufacturer’s protocols. No fragmentation or sizing was done on the samples before proceeding with ligation of Illumina-specific barcoded adapters and PCR amplification. The quality and concentration of the libraries were assessed using Nanodrop 2000 spectrophotometer (Thermo Scientific), the Qubit 3.0 dsDNA BR Assay (Thermo Scientific), D5000 ScreenTape (Agilent Technologies), and KAPA Library Quantification Kit (Roche Sequencing and Life Science), respectively, according to the manufacturers’ instructions.

Libraries were pooled and sequenced on the NovaSeq 6000 (Illumina) using the S4 Reagent Kit (Illumina) and100 bp paired-end reads. We targeted 30X coverage per sample, with the amount of generated data ranging from 33Gb to 112Gb across samples (see Supplementary Fig. S3).

### Evaluating museum-stored DNA for post-mortem damage

The long-term storage of museum specimens is associated with DNA degradation by deamination, leading to an excessive accumulation of cytosine to uracil (read by sequencer as thymine) changes ^27^. In downstream analyses, such post-mortem DNA damage can lead to biases and incorrect data interpretation. We evaluated the PAK genome sequences derived from the museum-stored samples using the Bayesian approach implemented in mapDamage 2.0, a program designed to track and quantify DNA damage patterns ^34^. Specifically, we focused our attention on the unusual accumulation of C to T and A to G mutations at the 5’ and 3’ termini as they represent the signatures of post-mortem deamination. Across the 14 PAK samples, the frequency of post-mortem DNA damage was estimated to be no greater than 0.1% of bases in each genome (Supplementary Fig. S2) and error signals were restricted to the 5bp within the 5’ and 3’ termini of reads. To eliminate potential biases and errors in our data, we trimmed the first and last 5 bases from each read using the “trimBam” option in “BamUtil” ^35^.

### Additional wild mouse genome sequences

We retrieved 155 previously published genomes belonging to *M. m. domesticus* (America, AMR = 50, France, FRA = 28, Germany, GER = 7, Iran = 7), *M. m. helgolandicus* (HEL, n = 3), *M. m. castaneus* (India, IND = 10, Taiwan, TAI = 20), *M. m. musculus* (Afghanistan, AFG = 6; Czech Republic, CZR = 8; Kazakhstan, KAZ = 8), and *M. spretus* (Spain, SPR = 8) ^24–26^. The newly sequenced 14 whole-genome data in this study were deposited to NCBI under SRA Project Accession Number PRJNA851025 (To be made public upon publication).

### Sequence alignment and variant calling

For the newly sequenced 14 PAK samples, we trimmed Illumina adapters using *cutadapt* ^36^. The clean reads were mapped to mm10 reference genome using the default parameters in BWA version 0.7.15 ^37^. Data from the 14 PAK samples were processed simultaneously with the 155 publicly available genome sequences to generate an integrated call set. We followed the standard Genome Analysis Toolkit (GATK; version 4.2) pipeline for subsequent pre-processing before variant calling ^38,39^. We performed variant calling for each sample separately using the “-ERC GVCF” mode in the “HaplotypeCaller”. Samples were then jointly genotyped using the “GenotypeGVCFs” GATK function and trained with previously ascertained mouse variants ^20^ using both the “VariantRecalibrator” and “ApplyVQSR” option of GATK. For the latter, the truth sensitivity level to initiate filtration was set to the default (i.e., 99). We filtered variants to exclude sites with missing alleles using VCFtools version 0.1.16 ^40^. All downstream analyses focus on autosomal bi-allelic single nucleotide variants.

### Analyses of population genetic structure

Principal component analysis was performed on all 169 wild mouse genomes using Plink version 2.0 ^41^. To construct a distance matrix tree, we first reformatted the variant file using “bcftools view file.vcf | bcftools query -f′%CHROM\t%POS[\t%TGT]\n′| sed -e ′s/\./N/g′” with BCFtools ^42^. We then used the python script “distMat.py” obtained from https://github.com/simonhmartin/genomics_general to generate the tree matrix ^43^. The tree was viewed using SplitsTree version 4.17.0 ^44^. We assessed the robustness of this topology to gene flow using TreeMix version 1.13 ^32^, allowing 0 – 5 migrations between any population pair in our dataset.

To calculate the allele sharing distance, we used the default option of “asd” (https://github.com/szpiech/asd) ^45^ and viewed the data using the R package “pheatmap” ^46^. We estimated the co-ancestry based Fst using the python program “popgenWindows.py” accessed from https://github.com/simonhmartin/genomics_general ^43^, and visualized results using the pheatmap R package ^46^.

### Inference of the dominant subspecies topology

We summarized the relationships among samples by building phylogenetic trees across 13,122 unique genomic regions, each defined by a fixed window of 50 SNPs. Trees were built for each window using the script “phyml_sliding_windows.py” accessed from https://github.com/simonhmartin/genomics_general. The output from the tree was used as input and weight assigned to each topology using *Twisst* - Topology Weighting by Iterative Sampling of Sub-Trees – based on the following options: --method complete, --abortCutoff 1000 --backupMethod fixed --iterations 400 ^47^. Topologies were viewed in R using the package “APE” version 5.5 ^48^.

## Data Availability

The raw fastq reads of the newly sequenced 14 wild house mice from Pakistan have been deposited in the NCBI Short Read Archive under the BioProject accession PRJNA851025 (https://www.ncbi.nlm.nih.gov/sra/PRJNA851025)

## Author contributions

RAL and BLD conceptualized the project. RAL performed all investigations and wrote the original draft with a major review by BLD. BLD was responsible for project supervision. Museum specimens were provided by VLM. AG generated the geographic map for all samples. MEB and JRC extracted the DNA and prepared genome sequencing libraries. All authors read, reviewed, and approved the final manuscript

---

**Supplementary Figure S1:**
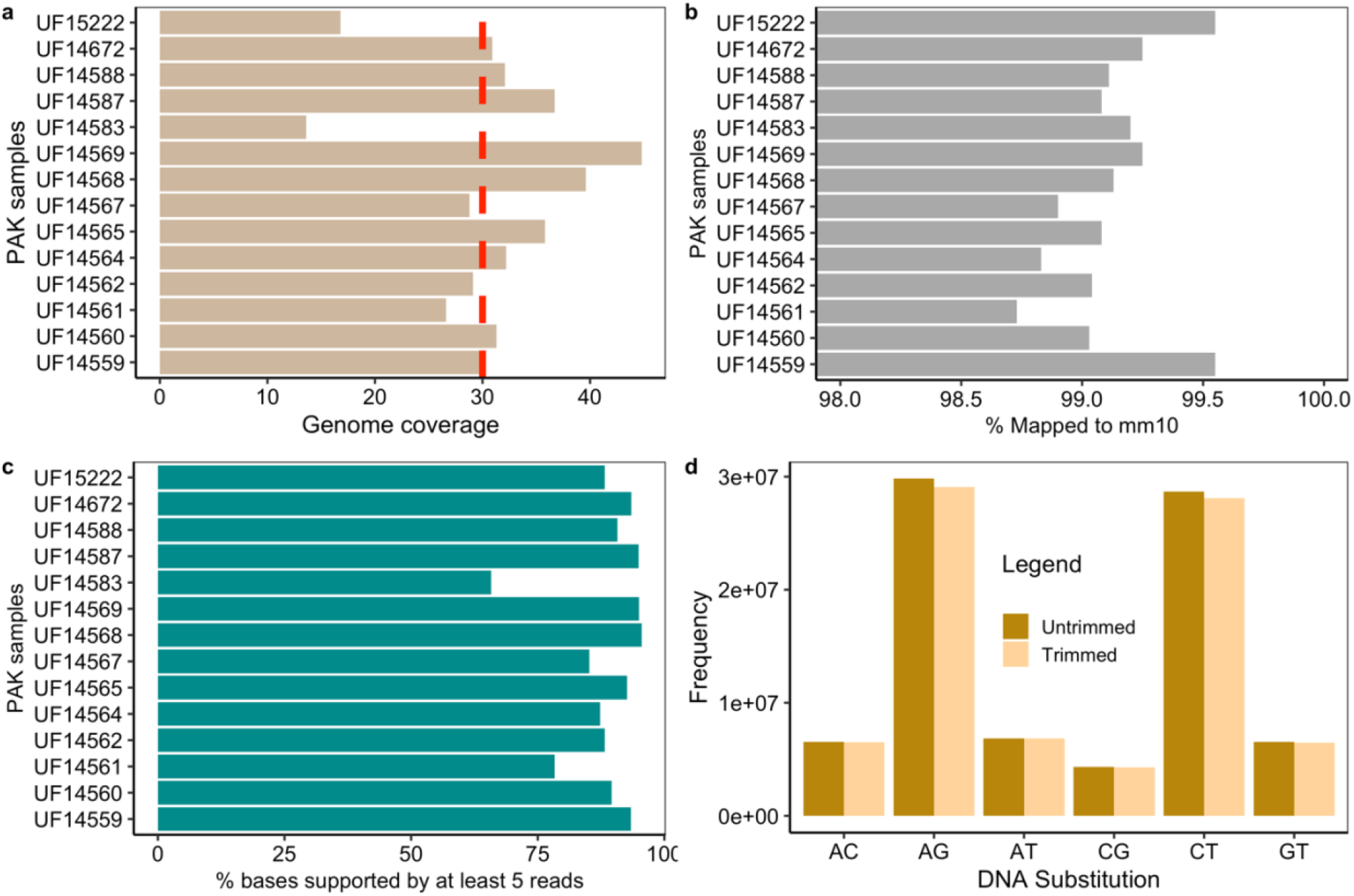
Sequence and mapping statistics for 14 PAK house mouse genomes. (**a**) We targeted an average of 30x coverage per sample (red line), with actual coverage values ranging from 17-45x. Bar charts showing (**b**) the percentage of sequenced reads mapping to the mm10 reference genome per sample and (**c**) the percentage of bases supported by at least 5 reads. (**d**) DNA substitution rates before and after trimming the 5bp at the 3’ and 5’ termini of each read.

**Supplementary Figure S2:**
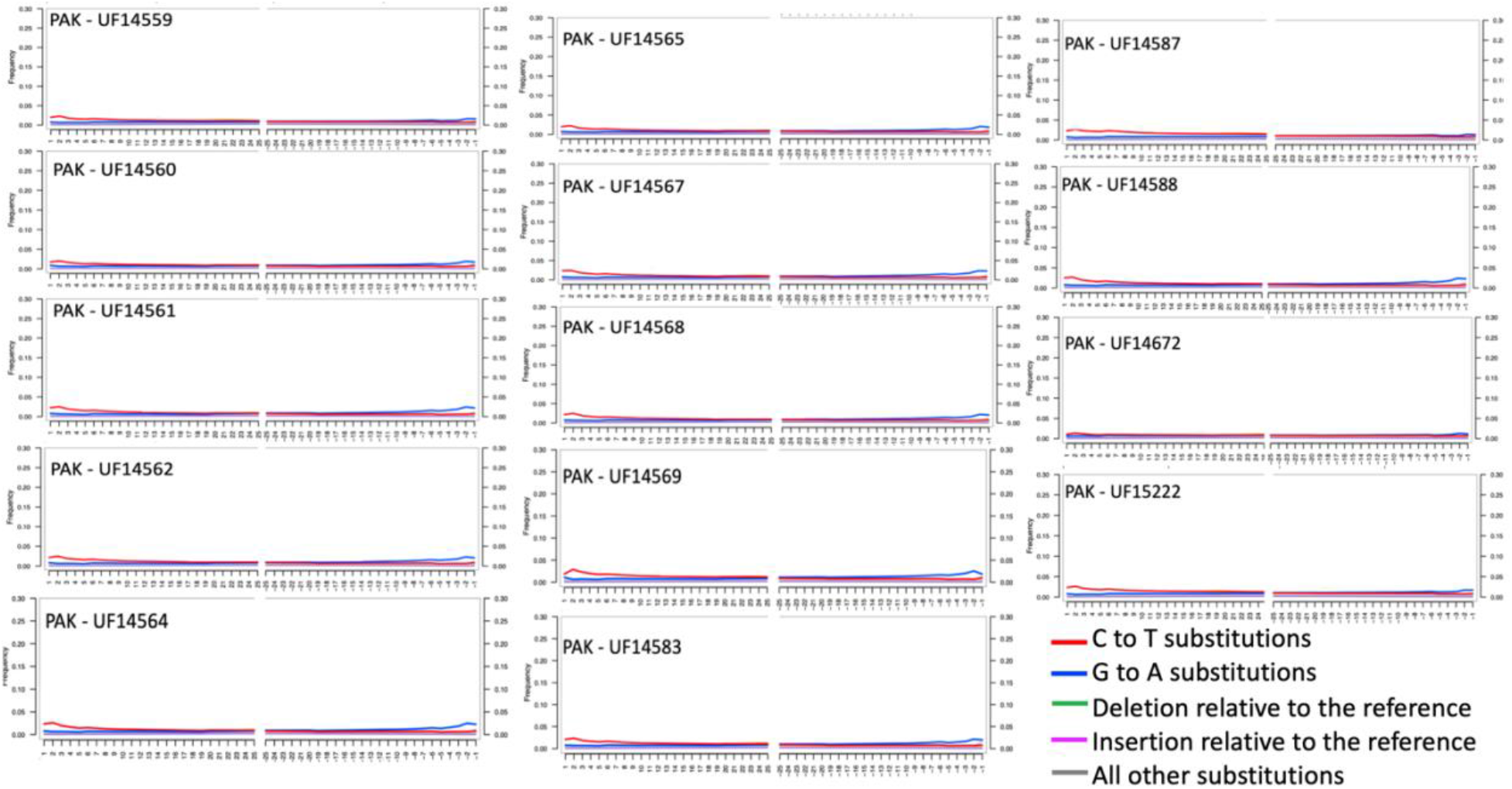
The distribution of DNA misincorporation errors at both the 5’ and 3’ termini of sequenced reads from PAK house mice. The y-axis shows the frequency of post-mortem DNA damage relative to the mm10 reference genome. Post-mortem DNA damage is expected to lead to an excess accumulation of C>T and G>A substitutions.

**Supplementary Figure S3:**
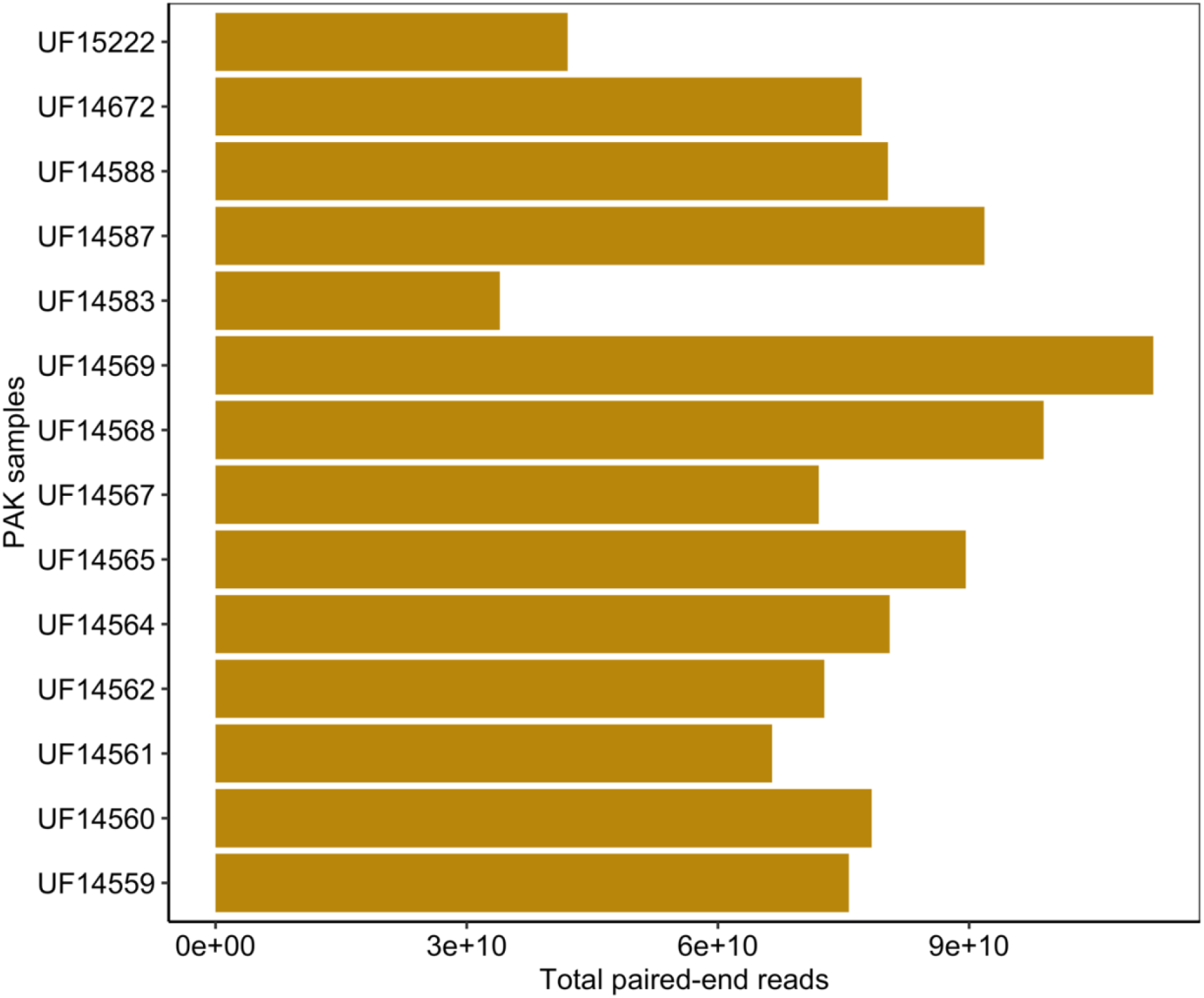
Bar plot of the number of paired-end reads generated for each PAK sample.

## References

1 Lawal, R. A., Arora, U. P. & Dumont, B. L. Selection shapes the landscape of functional variation in wild house mice. BMC Biol. 19, 1–17 (2021).

2 Suzuki, H., Shimada, T., Terashima, M., Tsuchiya, K. & Aplin, K. Temporal, spatial, and ecological modes of evolution of Eurasian Mus based on mitochondrial and nuclear gene sequences. Mol. Phylogen. Evol. 33, 626–646 (2004).

3 Suzuki, H., Aplin, K. & Pialek, J. Phylogeny and biogeography of the genus Mus in Eurasia. Evolution of the house mouse 3, 35 (2012).

4 Bonhomme, F. & Searle, J. B. House mouse phylogeography. Evolution of the house mouse 3, 278 (2012).

5 Suzuki, H. et al. Evolutionary and dispersal history of Eurasian house mice Mus musculus clarified by more extensive geographic sampling of mitochondrial DNA. Heredity 111, 375–390 (2013).

6 Geraldes, A., Basset, P., Smith, K. L. & Nachman, M. W. Higher differentiation among subspecies of the house mouse (Mus musculus) in genomic regions with low recombination. Mol. Ecol. 20, 4722–4736 (2011).

7 Boursot, P. et al. Origin and radiation of the house mouse: mitochondrial DNA phylogeny. J. Evol. Biol. 9, 391–415 (1996).

8 Mayr, E. Principles of systematic zoology. (Scientific Publishers, 2015).

9 Mayr, E. Animal species and evolution. Vol. 797 (Belknap Press of Harvard University Press Cambridge, Massachusetts, 1963).

10 Babiker, H. & Tautz, D. Molecular and phenotypic distinction of the very recently evolved insular subspecies Mus musculus helgolandicus ZIMMERMANN, 1953. BMC Evol. Biol. 15, 1–14 (2015).

11 Prager, E. M., Orrego, C. & Sage, R. D. Genetic variation and phylogeography of central Asian and other house mice, including a major new mitochondrial lineage in Yemen. Genetics 150, 835–861 (1998).

12 Boursot, P. et al. Origin and radiation of the house mouse: mitochondrial DNA phylogeny. J. Evol. Biol. 9, 391–415 (1996).

13 Duplantier, J.-M., Orth, A., Catalan, J. & Bonhomme, F. Evidence for a mitochondrial lineage originating from the Arabian peninsula in the Madagascar house mouse (Mus musculus). Heredity 89, 154–158 (2002).

14 Hardouin, E. A. et al. Eurasian house mouse (Mus musculus L.) differentiation at microsatellite loci identifies the Iranian plateau as a phylogeographic hotspot. BMC Evol. Biol. 15, 26 (2015).

15 Adhikari, P. et al. First molecular evidence of Mus musculus bactrianus in Nepal inferred from the mitochondrial DNA cytochrome B gene sequences. Mitochondrial DNA Part A 29, 561–566 (2018).

16 Zimmermann, K. Die hausmaus von helgoland. Mus musculus sspec Zeitschrift feur Seaugetierkunde 17, 163–166 (1953).

17 Yonekawa, H. et al. Evolutionary relationships among five subspecies of Mus musculus based on restriction enzyme cleavage patterns of mitochondrial DNA. Genetics 98, 801–816 (1981).

18 Yonekawa, H. et al. Hybrid origin of Japanese mice “Mus musculus molossinus”: evidence from restriction analysis of mitochondrial DNA. Mol. Biol. Evol. 5, 63–78 (1988).

19 Schoch, C. L. et al. NCBI Taxonomy: a comprehensive update on curation, resources and tools. Database 2020 (2020).

20 Keane, T. M. et al. Mouse genomic variation and its effect on phenotypes and gene regulation. Nature 477, 289 (2011).

21 White, M. A., Ané, C., Dewey, C. N., Larget, B. R. & Payseur, B. A. Fine-scale phylogenetic discordance across the house mouse genome. PLoS Genet. 5, e1000729 (2009).

22 Phifer-Rixey, M., Harr, B. & Hey, J. Further resolution of the house mouse (Mus musculus) phylogeny by integration over isolation-with-migration histories. BMC Evol. Biol. 20, 1–9 (2020).

23 Yang, H. et al. Subspecific origin and haplotype diversity in the laboratory mouse. Nat. Genet. 43, 648 (2011).

24 Harr, B. et al. Genomic resources for wild populations of the house mouse, Mus musculus and its close relative Mus spretus. Sci Data 3, 160075 (2016).

25 Phifer-Rixey, M. et al. The genomic basis of environmental adaptation in house mice. PLoS Genet. 14, e1007672 (2018).

26 Davies, R. W. Factors influencing genetic variation in wild mice, University of Oxford, (2015).

27 Sawyer, S., Krause, J., Guschanski, K., Savolainen, V. & Pääbo, S. Temporal patterns of nucleotide misincorporations and DNA fragmentation in ancient DNA. PloS One 7, e34131 (2012).

28 Kirch, M., Romundset, A., Gilbert, M. T. P., Jones, F. C. & Foote, A. D. Ancient and modern stickleback genomes reveal the demographic constraints on adaptation. Curr. Biol. 31, 2027–2036. e2028 (2021).

29 Briggs, A. W. et al. Removal of deaminated cytosines and detection of in vivo methylation in ancient DNA. Nucleic Acids Res. 38, e87–e87 (2010).

30 GBIF.org (25 April 2022) GBIF occurrence download (https://doi.org/10.15468/dl.xuksm3).

31 Teeter, K. C. et al. Genome-wide patterns of gene flow across a house mouse hybrid zone. Genome Res. 18, 67–76 (2008).

32 Pickrell, J. K. & Pritchard, J. K. Inference of population splits and mixtures from genome-wide allele frequency data. PLoS Genet. 8, e1002967, doi:https://doi.org/10.1371/journal.pgen.1002967 (2012).

33 Moraes-Barros, N. d. & Morgante, J. S. A simple protocol for the extraction and sequence analysis of DNA from study skin of museum collections. Genetics and Molecular Biology 30, 1181–1185 (2007).

34 Jónsson, H., Ginolhac, A., Schubert, M., Johnson, P. L. & Orlando, L. mapDamage2. 0: fast approximate Bayesian estimates of ancient DNA damage parameters. Bioinformatics 29, 1682–1684 (2013).

35 Jun, G., Wing, M. K., Abecasis, G. R. & Kang, H. M. An efficient and scalable analysis framework for variant extraction and refinement from population-scale DNA sequence data. Genome Res. 25, 918–925 (2015).

36 Martin, M. Cutadapt removes adapter sequences from high-throughput sequencing reads. EMBnet. journal 17, 10–12 (2011).

37 Li, H. Aligning sequence reads, clone sequences and assembly contigs with BWA-MEM. arXiv preprint arXiv:1303.3997 (2013).

38 Li, H. & Durbin, R. Fast and accurate long-read alignment with Burrows–Wheeler transform. Bioinformatics 26, 589–595, doi:10.1093/bioinformatics/btp698 (2010).

39 Auwera, G. A. et al. From FastQ data to high-confidence variant calls: The genome analysis toolkit best practices pipeline. Curr Protoc Bioinform 43, 11.10.11–33, doi:10.1002/0471250953.bi1110s43 (2013).

40 Danecek, P. et al. The variant call format and VCFtools. Bioinformatics 27, 2156–2158, doi:10.1093/bioinformatics/btr330 (2011).

41 Chang, C. C. et al. Second-generation PLINK: rising to the challenge of larger and richer datasets. Gigascience 4, 7, doi:https://doi.org/10.1186/s13742-015-0047-8 (2015).

42 Danecek, P. et al. Twelve years of SAMtools and BCFtools. Gigascience 10, giab008 (2021).

43 Martin, S. H., Davey, J. W. & Jiggins, C. D. Evaluating the use of ABBA–BABA statistics to locate introgressed loci. Mol. Biol. Evol. 32, 244–257, doi:10.1093/molbev/msu269 (2015).

44 Huson, D. H. & Bryant, D. Application of phylogenetic networks in evolutionary studies. Mol. Biol. Evol. 23, 254–267 (2006).

45 Gao, X. & Starmer, J. Human population structure detection via multilocus genotype clustering. BMC Genet. 8, 1–11 (2007).

46 Kolde, R. Pheatmap: pretty heatmaps. R package version 1, 747 (2012).

47 Martin, S. H. & Van Belleghem, S. M. Exploring evolutionary relationships across the genome using topology weighting. Genetics 205 (2017).

48 Paradis, E., Schliep, K. & Schwartz, R. APE 5.0: an environment for modern phylogenetics and evolutionary analyses in R. Bioinformatics 1, 3, doi:https://doi.org/10.1093/bioinformatics/bty633 (2018).

